# Microprotein miP-PSTPIP2 affects cytoskeleton dynamics to modulate endothelial cell endocytosis, barrier function and migration

**DOI:** 10.1101/2025.11.07.687176

**Authors:** Beyza Güven, Bethlehem Bezuneh, Manav Raheja, Witold Szymanski, Ralf P. Brandes, Johannes Graumann, Ingrid Fleming, Mauro Siragusa

**Affiliations:** Goethe University, Institute for Vascular Signalling, Centre for Molecular Medicine, Frankfurt am Main, Germany; Philipps-Universität Marburg, Institute of Translational Proteomics & Core Facility Translational Proteomics, Biochemical/Pharmacological Centre, Marburg, Germany; Institute for Cardiovascular Physiology, Goethe-University, Frankfurt, Germany; German Center for Cardiovascular Research (DZHK), Partner site RheinMain, Frankfurt am Main and Bad Nauheim, Germany; CardioPulmonary Institute, Frankfurt am Main and Bad Nauheim, Germany

**Keywords:** small open reading frames, microproteins, endothelial cell, endocytosis, cardiovascular disease

## Abstract

**Background:** A large number of microproteins (miPs) encoded by small open reading frames (smORFs) are expressed in endothelial cells, yet their function remains largely unknown. In this study, we characterized a novel 46-amino-acid miP encoded by a smORF within the proline-serine-threonine phosphatase interacting protein 2 (*PSTPIP2*) transcript that was upregulated under inflammatory conditions and we refer to as miP-PSTPIP2.

**Methods:** Immunoprecipitation coupled with mass spectrometry-based proteomics, immunoblotting, immunofluorescence and proximity ligation assays were used to identify and validate miP-PSTPIP2 interacting proteins in human endothelial cells. The impact of adenovirus-mediated overexpression of miP-PSTPIP2 on endocytosis, cytoskeleton dynamics and abundance of proteins involved in these processes was investigated by confocal microscopy and immunoblotting. Live cell imaging was used to assess endothelial cell migration and vascular permeability.

**Results:** miP-PSTPIP2 physically associated with caveolar proteins, proteins involved in the regulation of cytoskeleton dynamics, intracellular transport, clathrin adaptor activity, as well as nuclear proteins. Human endothelial cells overexpressing miP-PSTPIP2 demonstrated enhanced endocytosis and transcytosis of transferrin as well as low-density lipoprotein. Mechanistically, miP-PSTPIP2 modulated Arp2/3-mediated actin nucleation and branching, which are required for dynamic cytoskeleton rearrangements. Moreover, altered cytoskeleton dynamics in miP-PSTPIP2-expressing endothelial cells resulted in impaired cell migration as well as increased permeability and monocyte trans-endothelial migration.

**Conclusions:** miP-PSTPIP2 is an inflammation-induced endothelial miP that regulates Arp2/3-dependent actin dynamics, thereby enhancing lipid uptake and leukocyte permeability. Its upregulation under inflammatory conditions suggests a contributory role in endothelial dysfunction and vascular inflammation.

## INTRODUCTION

The realization that the translation of non-canonical small open reading frames (smORFs) can result in the generation of microproteins (miPs) is relatively recent.^1–7^ Because of their size i.e. fewer than 100 amino acids, challenges in their detection have limited their characterization. Despite this, miPs are already known to be involved in diverse biological processes including cell growth and metabolic regulation, with potential implications in human disease. MiPs often exert their function via the formation of macromolecular complexes to participate in cell signaling and the modulation of crucial cellular processes.^8–10^ Numerous miPs have been reported to play a role in the heart and heart disease by modulating cardiac contractility, mitochondrial energy production and calcium handling.^11–16^ However, the extent to which miPs influence endothelial cell function and vascular homeostasis is largely unknown.

Endothelial cells form the inner lining of blood vessels and play an essential role in the modulation of vascular homeostasis, inflammation, and vascular repair. To-date, ribosome profiling and proteogenomic approaches have demonstrated that thousands of smORFs are translated in endothelial cells and that their expression is altered under inflammatory conditions.^17–19^ We found that the expression of hundreds of endothelial cell smORFs was altered in a mouse model of endothelial dysfunction and accelerated atherogenesis as well as in human endothelial cells treated with interleukin-1β (IL-1β).^19^ Among the human smORFs with a mouse homolog that were upregulated under inflammatory conditions *in vivo* and *in vitro*, we identified a smORF located within the proline-serine-threonine phosphatase interacting protein 2 (*PSTPIP2*) transcript (ENST00000409746.5 in human and ENSMUST00000114741.4 in mouse), albeit in a different reading frame than the canonical coding sequence. The smORF encodes a 46-amino acid miP in humans and a 54-amino acid miP in mice, referred to as miP-PSTPIP2. We have generated and validated a custom antibody against this miP. Using this antibody, we demonstrated that the miP localizes to the actin cytoskeleton and in proximity to the plasma membrane, within cytosolic vesicles and in the nucleus. Consistent with the smORF translatome data, miP-PSTPIP2 expression was upregulated under inflammatory conditions *in vitro* and *in vivo*, and its presence was also detected in the endothelial layer of human carotid atherosclerotic plaques.^19^ The aim of this study was to characterize the mechanism of action of endothelial cell miP-PSTPIP2.

## METHODS

### Cell culture

Human umbilical vein endothelial cells were isolated and cultured as described previously^20,21^ and used up to passage 4. The use of human material in this study complies with the principles outlined in the Declaration of Helsinki (World Medical Association, 2013), and the isolation of endothelial cells was approved in written form by the ethics committee of the Goethe-University. THP-1 monocytic cells were obtained from the American Type Culture Collection (LGC Standards, Lancashire, United Kingdom) and cultured in RPMI-1640 containing 2 mmol/L glutamine, 10 mmol/L HEPES, 1 mmol/L sodium pyruvate, 4.5g/L glucose, 1.5 g/L sodium bicarbonate and 10% fetal bovine serum (FBS). Porcine aortic endothelial cell (PAEC) were kindly provided by Dr. J. Waltenberger (Maastricht, The Netherlands), cultured in F-12 Nut Mix (Ham) containing 1.5 g/L sodium bicarbonate and 8% FBS and used up to passage 9. All cells were negative for mycoplasma contamination. Cultured cells were kept in a humidified incubator at 37°C containing 5% CO_2_.

### Generation of adenoviruses

The adenoviral vector used to overexpress FLAG-tagged miP-PSTPIP2 was constructed and packaged by VectorBuilder (Chicago, United States). Briefly, the sequence encoding the FLAG tag (GACTACAAAGACGATGACGACAAG) was inserted on the 5’ end of the human smORF-PSTPIP2 sequence (GGCAGAACAGGCCGTCAGCCGGAGTGCCAACCTGGTGAACCCGAAGCAACAA GAAAAGCTTTTTGTGAAACTGGCAACTTCAAAGACCGCAGTAGAGGACTCAGACAAAGCATACATGCTGCACATCGGCACCCTGGA), preceded by Kozak sequence (GCCACC) and ATG start codon and followed by the stop codon TAA. The restriction sites AbsI and SgrDI were placed on the 5’ and 3’ end on of the insert, respectively (VectorBuilder ID: VB230219-1395jcp). The insert was cloned into the mammalian gene expression adenoviral vector pAd5 under the CMV promoter. DNA sequencing confirmed correct direction of the insert and lack of mutations. Replication incompetent adenoviruses were generated by transfection into packaging cells. The adenoviral vector and replication incompetent adenoviruses used to overexpress EGFP, which served as a control, were also constructed and packaged by VectorBuilder as described above (VectorBuilder ID: VB010000-9299hac).

### Adenoviral transduction and treatment of endothelial cells

#### FLAG-miP-PSTPIP2/EGFP

Human endothelial cells (passage 4, 90% confluent) were starved of serum in endothelial cell basal medium (EBM, PromoCell, Heidelberg, Germany) containing 0.1% BSA and transduced overnight with adenoviruses to express the FLAG-miP-PSTPIP2 or EGFP as control (250 MOI). For functional assays, the medium was replaced after 24 hours to endothelial cell growth medium 2 (ECGM2, PromoCell) containing 2% FBS and cells were used 48 hours after infection.

### Cell lysis and miP-PSTPIP2 immunoprecipitation

Three independent cell batches of human endothelial cells (passage 4) were grown in ∅10 cm dishes until confluent and then treated with human recombinant IL-1β (20 ng/mL, Peprotech, Hamburg, Germany) for 20 hours. Thereafter, cells were washed with phosphate-buffered saline (PBS) and lysed in a buffer (1 mL) containing Tris/HCl pH 7.5 (50 mmol/L), NaCl (150 mmol/L), NP-40 (1%), NaPPi (10 mmol/L), NaF (20 mmol/L), orthovanadate (2 mmol/L), okadaic acid (10 nmol/L), β-glycerophosphate (50 mmol/L), phenylmethylsulfonyl fluoride (230 µmol/L) and an EDTA-free protease inhibitor mix (AppliChem GmbH, Darmstadt, Germany). Lysates were incubated for 45-60 minutes at 4°C on an end-over-end rocker and gently vortexed until no cell clumps were visible. Protein concentration was determined using the Bradford method. Whole cell lysates (1.2 mg/sample) were incubated overnight with anti-miP-PSTPIP2 (6 µg/mg, Eurogentec) or control rabbit IgGs (Cat. Nr.: NI01, Merck) in an end-over-end rocker at 4°C. This was followed by the addition of packed rec-Protein G-Sepharose 4B Conjugate (50 µL/mg lysate, Invitrogen) for 2 hours, again under constant agitation 4°C. Samples were then centrifuged (2200 g, 2 minutes) and the supernatant removed. The remaining pellet was washed twice with affinity purification lysis buffer and centrifuged (2200 g, 30 seconds). Finally, pellets were washed with a buffer containing Tris HCl pH 7.5 (50 mmol/L) and NaCl (150 mmol/L), centrifuged (2200 g, 30 seconds, 4°C) before the supernatant was carefully removed. The dry pellets were frozen at -20°C until further processing for proteomic analyses. For immunoblotting, samples were boiled in elution buffer (2% SDS, 1% β-mercaptoethanol and 0.005% bromophenol blue in PBS) for 5 min at 95°C.

### Identification of the miP-PSTPIP2 interactome by LC-MS/MS

Immunoprecipitates were resuspended in 20 μL of 8% sodium lauroyl sarcosinate (SLS) in 100 mmol/L Triethylammonium bicarbonate solution (pH∼7.5) and boiled for 10 minutes at 90°C in a ThermoMixer (1800 rpm). Samples were reduced by incubating with dithiothreitol (1 μL of 400 mmol/L stock to give a final concentration of 10 mmol/L) for 10 minutes (95°C). This was followed by alkylation (30 minutes, 25°C) using iodoacetamide (1 μL of 550 mmol/L stock to give a final concentration of 13 mmol/L). All samples were diluted with 50 mmol/L TEAB buffer to a final volume of 400 μL and trypsin was added (final amount 1 μg). Digestion was allowed to continue for 16 hours (37°C) and was stopped by addition of trifluoroacetic acid (final concentration 1.5 %). Precipitating lauroyl sarcosinate was removed by centrifugation and peptides were purified using solid phase extraction on 2 discs of C18 StageTips.^22^ Purified peptides were first dried, then resuspended in 40 μL of 0.1% trifluoroacetic acid. Peptide concentrations were determined using the Pierce Fluorimetric Peptide Assay, and sample volumes were adjusted in order to allow injection of equal amounts of peptides onto the analytical column. Subsequently, 500 ng of peptides were loaded onto a µPAC™ -Trapping-column (2 mm diameter, 10 mm length) and eluted with a gradient from 98% solvent A (0.1% formic acid) and 2% solvent B (99.9% acetonitrile and 0.1% formic acid) to 7% solvent B over 15 minutes, continued from 7 to 38% of solvent B for another 90 minutes, then from 38 to 55% of solvent B for 15 minutes over a reverse-phase high-performance liquid chromatography (HPLC) separation column (Thermo Scientific™ µPAC™ HPLC Columns, 50 cm) with a flow rate of 300 nL/minutes. Peptides were infused via an Advion TriVersa NanoMate (Advion BioSciences, Inc. New York, United States) into an Orbitrap Eclipse Tribrid mass spectrometer (Thermo Scientific). The mass spectrometer was operating in positive-ionization mode with a spray voltage of the NanoMate system set to 1.7 kV and source temperature at 275°C. Data were acquired by means of a data-independent acquisition paradigm. In short, spectra were acquired with a MS1 resolution of 60,000 and mass range from 375 to 1,500 m/z for the precursor ion spectra, maximum injection time was set to 50 milliseconds, Automatic Gain Control target of 400k and normalized AGC target of 100%. This was followed by data-independent acquisition scans with 40 fixed windows of 15 m/z width and 1 m/z of overlap, ranging from 375 to 975 m/z. MS2 spectra were acquired with the resolution of 30,000, collision energy of 30%, maximum injection time of 54 milliseconds and AGC target of 2000%. Peptide spectrum matching and label-free quantitation were subsequently performed using DIA-NN^23^ with a library-free search against the Human https://Uniprot.org database (20388 reviewed Swiss-Prot entries; September 2022). In brief, output was filtered to a 0.01 FDR on the precursor level. Deep learning was used to generate an in silico spectral library for library-free search. Fragment m/z was set to a minimum of 200 and a maximum of 1,800. In silico peptide generation allowed for N-terminal methionine excision, tryptic cleavage following K*,R*, a maximum of one missed cleavage, as well as a peptide length requirement of seven amino acid minimum, and a maximum of 30. Cysteine carbamidomethylation was included as a fixed modification and methionine oxidation (maximum of two) as a variable modification. Precursor masses from 300 to 1,800 m/z and charge states one to four were considered. DIA-NN was instructed to optimize mass accuracy separately for each acquisition analysed and protein sample matrices were filtered using a run-specific protein q-value (“--matrix-spec-q” option). Log files from DIA-NN processing can be found in the ProteomeXchange repository. Downstream data processing and statistical analysis were carried out by the the limma-based Autonomics package developed in-house (DOI: 10.18129/B9.bioc.autonomics; version 1.1.7.6).^24^ Proteins with a q-value of <0.01 were included for further analysis. MaxLFQ^25^ values were used for quantitation and missing values were imputed. DIA-NN initially identified 5813 protein groups. All intensities and maxLFQ values containing only 1 precursor (Np) per sample were replaced with NA in that particular sample.

### Interactome analysis

For the miP-PSTPIP2 interactome, proteins significantly (P value ≤0.05) enriched in the miP-PSTPIP2 pulldown compared to the control IgG pulldown were analyzed by STRING v.11.5 GO term enrichment analysis.^26^

### Immunoblotting

Protein samples were separated by SDS–PAGE and transferred to 0.45 µm nitrocellulose membranes (GE Healthcare, Freiburg, Germany). Membranes were incubated overnight with the following primary antibodies: AP2A1 (1:1000, Novus Biologics, Centennial, United States, NB600-1545), AP2M1 (1:1000, Cell Signalling, Danver, United States, #68196), Arp2 (1:2000, Proteintech, Rosemont, United States, 10922-1-AP), Arp3 (1:1000, Abcam, Cambridge, United Kingdom, ab49671), β-actin (1:1000, Sigma-Aldrich, Burlington, United States, A1978), CAV1 (1:1000, Cell Signaling, #3238), CHC (1:2000, BD Transduction, Franklin Lakes, United States, 610499), Cortactin (1:10000, Abcam, ab81208), DAAM1 (1:2000, Proteintech, 67287-1-Ig), GAPDH (1:300, Millipore, Burlington, United States, MAB374), LDLR (1:2000, Invitrogen, Waltham, United States, PA5-46985), Phospho-AP2M1 (1:1000, Cell Signaling, #7399), RhoA (1:1000, Santa Cruz, Dallas, United States, sc-179) and VASP (1:1000, BD Transduction, V40020). Thereafter, membranes were incubated with species-specific secondary antibodies anti-IgG conjugated with horseradish peroxidase. Proteins were visualized by enhanced chemiluminescence using a commercially available kit (GE Healthcare, Chicago, United States).

### Immunofluorescence

Cells grown on 8-well chamber slides (Ibidi, Martinsried, Germany) were washed with PBS supplemented with 0.8 mmol/L CaCl₂ and 1.4 mmol/L MgCl₂ and fixed in 4% paraformaldehyde (Carl Roth, Karlsruhe, Germany). Cells were then incubated for 30 minutes at room temperature in blocking/permeabilization buffer containing 5% horse serum and 0.1% Triton X-100 in PBS. Primary antibodies against AP2A1 (1:100, Novus Biologics, NB600-1545), miP-PSTPIP2 (5 µg/ml, Eurogentec, Seraing, Belgium)^19^, control rabbit IgG (5 µg/ml, Millipore, NI01) or phalloidin-Alexa Fluor 633 (1:300, Invitrogen, A22284) were diluted in PBS containing 0.5% horse serum and 0.01% Triton X-100, and incubated for 2 hours at room temperature. Cells were subsequently incubated for 1 hour at room temperature with appropriate species-specific donkey Alexa Fluor-conjugated secondary antibodies (Thermo Fisher Scientific, Waltham, United States). Nuclei were stained with DAPI (Thermo Fisher Scientific). Slides were mounted in mounting medium (43% glycerol, 100 mmol/L DTT). Images were acquired with a confocal microscope (STELLARIS 8 FALCON; Leica, Wetzlar, Germany) and processed using LAS AF lite software (Leica).

### Proximity ligation assay

Cells were fixed as described above and permeabilized with 0.1% Triton X-100 and blocked using the blocking solution provided in the Duolink In Situ kit (Sigma-Aldrich, DUO92002, DUO92004, DUO92008). Primary antibodies were applied for 2 h at room temperature: miP-PSTPIP2 (5 µg/ml, Eurogentec), Arp3 (1:1000, Abcam, ab49671), CAV1 (1:100, Transduction Laboratories, C43420) and CHC (1:2000, BD Transduction, 610499). The proximity ligation assay was performed using the Duolink In Situ Detection reagents according to the manufacturer’s instructions. Nuclei were stained with DAPI (Thermo Fisher Scientific), and cells were co-stained with VE-cadherin (1:200, R&D Systems, Minneapolis, United States, AF938) and donkey anti-goat Alexa Fluor647-conjugated secondary antibody (Thermo Fisher Scientific) or phalloidin-Alexa Fluor 633 (1:400, Invitrogen, A22284) where indicated. Slides were mounted in mounting medium (43% glycerol, 100 mmol/L DTT). Images were acquired with a confocal microscope (STELLARIS 8 FALCON; Leica) and processed using LAS AF lite software (Leica).

### Functional assays

#### Endocytosis

Human or porcine endothelial cells were seeded at 60% confluency in ibiTreat µ-Slide 8-well chambers (ibidi). Following overnight incubation in EBM medium supplemented with 0.1% BSA, cells were incubated at 37°C with ECGM2 medium containing either 25 µg/mL Alexa Fluor 555-labeled transferrin (Cat. No.: T35352, Invitrogen) for 5, 10 or 15 minutes, or 10 µg/mL DiI-labeled LDL (Cat. No.: L3482, Invitrogen), Dil-oxLDL (Cat. No.: L34358, Invitrogen) or Dil-acLDL (Cat. No.: L3484, Invitrogen) for 60 minutes. Cells were then washed twice with ice-cold PBS and fixed with 4% Rotifix (Carl Roth) for 15 minutes at room temperature. Fluorescent images were acquired using a confocal microscope (STELLARIS 8 FALCON; Leica) and LAS AF lite software (Leica) and mean fluorescence intensity (MFI) was quantified determine transferrin or LDL uptake.

#### Transcytosis

Human endothelial cells were seeded at 100% confluency in fibronectin-coated Transwell cell culture inserts (1.0 µm pore size, Sarstedt, Nümbrecht, Germany, Cat. No.: 83.3932.101) placed in 24-well plates. After an additional 24 hours, cells were incubated in EBM supplemented with 0.1% BSA overnight. Inserts were incubated with 40 µg/mL Alexa Fluor 555-labeled transferrin (Cat. No.: T35352, Invitrogen) or 25 µg/mL DiI-labeled LDL (Cat. No.: L3482, Invitrogen) in EBM + 0.1% BSA for 60 minutes at 37°C. Cells were subsequently washed three times with EBM + 0.1% BSA, and inserts were transferred to fresh wells containing ECGM2 medium to allow transcytosed material to accumulate in the lower compartment for an additional 90 minutes. Medium from the lower compartment was collected and MFI was measured using a PerkinElmer 2104 EnVision Multilabel Plate Reader (Waltham, United States).

#### Permeability

Cells were seeded at 100% confluency in fibronectin-coated transwell cell culture inserts (1.0 µm pore size, Sarstedt, Cat. No.: 83.3932.101) placed in 24-well plates. After 24 hours, cells were starved of serum in phenol red-free DMEM/F12 medium supplemented with 0.1% BSA overnight. The medium in both compartments was replaced and FITC-conjugated 40 kDa dextran (Sigma-Aldrich, Cat. No.: FD40) was added to the upper compartment at a final concentration of 1 mg/mL and incubated for 15 minutes at 37°C. The medium from the lower compartments was then collected to measure the amount of FITC-conjugated 40 kDa dextran. Samples were transferred to a 96-well UV-transparent plate, along with blank (medium) and a standard curve prepared with increasing amounts of FITC-dextran and MFI was measured using a PerkinElmer EnVision Multilabel Plate Reader (PerkinElmer).

#### Transendothelial migration

Cells were seeded in fibronectin-coated Transwell inserts (8.0 µm pore size, Sarstedt, Cat. No.: 83.3932.800) at a density of approximately 100,000 cells/insert. Inserts were placed into 24-well plates, with 200 µL of ECGM2 medium added to the apical (insert) chamber and 1 mL to the basolateral (lower) compartment. Cells were cultured for 72 hours. THP-1 monocytes were stained using CellTracker Red CMTPX (Thermo Fisher Scientific) at a final concentration of 200 nmol/L. Briefly, cells were resuspended in ECGM2 medium and incubated with dye for 15 minutes at 37°C. Stained cells were washed, counted, and resuspended in ECGM2 medium for seeding at 25,000 cells per insert. Migration was stimulated by adding ECGM2 medium supplemented with 50 ng/mL monocyte chemotactic protein-1 (R&D Systems, Cat. No.: 279-MC) to the lower compartment. Transendothelial migration was monitored using the IncuCyte S3 Live-Cell Analysis System (Essen BioScience, Ann Arbor, United States) with 20× objective, capturing phase contrast and red fluorescence channels across 36 fields per well for up to 6 hours. Image analysis of red-labeled THP-1 cells in the lower chamber was performed using the following parameters: segmentation method = AI Confluence; radius = 100 µm; hole fill = 0 µm²; size adjustment = 0; area filter = minimum 7 µm².

#### Scratch wound

Cells (40,000 cells per well) were seeded in ImageLock 96-well cell culture plates (Sartorius, Cat. No.: BA-04855) in ECGM2 medium (PromoCell). After 24 hours, confluent monolayers were scratched across the well center using a plastic pipette tip, and wells were washed once with medium to remove detached cells. Wound closure was monitored for up to 48 hours using the IncuCyte S3 Live-Cell Analysis System (Essen BioScience) equipped with a 10× objective, capturing and analyzing phase-contrast images using the Scratch Wound assay module.

#### Directed migration

Cells (4,000 cells per well) were seeded in the central reservoir of an ibidi µ-Slide Chemotaxis (ibidi, Cat. No.: 80326). After 2 hours of attachment, EBM medium supplemented with 0.1% BSA was added to both reservoirs, with the right reservoir further supplemented with 10% FBS to establish a chemotactic gradient. Cell migration was imaged over 20 hours at 10-minute intervals using a Zeiss Axio Observer inverted microscope equipped with an 10x objective (Zeiss, Oberkochen, Germany). Images were acquired using ZEN Blue 3.3 software (Zeiss). Cell tracks were manually annotated with the ImageJ Manual Tracking plugin, and migration parameters were analyzed using the Chemotaxis and Migration Tool v2.0 (ibidi). Only cells migrating toward the FBS gradient were included for quantification.

#### Actin nucleation

Cells were treated with 0.5 µmol/L cytochalasin D (CytoD, AppliChem, Darmstadt, Germany, A7641-0001) for 20 minutes to disrupt the actin cytoskeleton. Cells were subsequently incubated in ECGM2, fixed with 4% paraformaldehyde (Rotifix, Carl Roth) immediately or after the indicated time points (up to 180 minutes after treatment) and processed for immunofluorescence staining. All samples were imaged using a STELLARIS confocal microscope (Leica) equipped with a 63x oil immersion objective. Full z-stacks were acquired for each condition.

### Total Internal Reflection Fluorescence microscopy

Cells were stimulated with 25 µg/mL Alexa Fluor 555–conjugated transferrin (Invitrogen, T35352) in ECGM2 medium for 1, 5 or 15 minutes at 37 °C and fixed in 4% paraformaldehyde (Rotifix, Carl Roth) for 15 minutes at room temperature. Cells were then stained for miP-PSTPIP2 as described above. Total Internal Reflection Fluorescence imaging was performed using a Zeiss Axio Observer inverted microscope equipped with an α Plan-Apochromat 100×/1.46 oil immersion objective (Zeiss). Images were acquired using Zen Blue 3.3 software (Zeiss) and processed identically to confocal images.

### Flow cytometry

Cells were pelleted and resuspended in PBS containing 0.5% BSA on ice. Following an additional wash in PBS containing 0.5% BSA, cells were incubated with BD OptiBuild™ RB705 mouse anti-human LDLR antibody (BD Biosciences, Cat. No.: 757511) at a 1:100 dilution for 30 minutes on ice in the dark. Cells were then fixed with 4% paraformaldehyde (Rotifix, Carl Roth), washed once with PBS containing 0.5% BSA, and resuspended in PBS containing 0.1% BSA. Fluorescence was analyzed on a BD LSR II/Fortessa flow cytometer (BD Biosciences, Heidelberg, Germany), and data were processed using FlowJo Vx software (TreeStar, Ashland, United States). Daily instrument calibration was performed with Cytometer Setup and Tracking beads (BD Biosciences).

### AlphaFold prediction of structure

Protein structure predictions were obtained using AlphaFold-3 (DeepMind / Google) via the AlphaFold Server.^27^ The input protein amino acid sequence was provided in FASTA format, with no template restrictions. Structural models were visualized using PyMOL (The PyMOL Molecular Graphics System, Version 3.0 Schrödinger, LLC).

### Statistics

Results are presented as mean ± SEM. GraphPad Prism software (v. 10.1.2) was used to assess statistical significance. Differences between two groups were compared by two-tailed unpaired Student’s t-test. All experiments in which the effects of two variables were tested were analyzed by two-way ANOVA followed by the Holm-Šídák’s multiple comparisons test. Differences were considered statistically significant when P < 0.05. Only exact significant P values are reported.

### Data and materials availability

Further information and requests for resources and reagents should be directed to and will be fulfilled by Mauro Siragusa. Data that support the findings of this study are available as part of the manuscript. Raw data can be accessed at the following repositories:

Mass spectrometry-based proteomics data: ProteomeXchange Consortium via the MassIVE partner repositories (http://proteomecentral.proteomexchange.org) with the following identifiers:

miP-PSTPIP2 interactome: PXD050588 (MassIVE ID: MSV000094309)

Reviewer account details:

Username: MSV000094309_reviewer

Password: ipmips_001_2024

## RESULTS

### miP-PSTPIP2 associates with proteins related to intracellular transport

The smORF-PSTPIP2 spans exons 7 to 9 of the *PSTPIP2* gene, nested within the coding sequence (CDS) in the +2 reading frame. Structure prediction (AlphaFold 3^27,28^) suggested that miP-PSTPIP2 consists of two alpha-helical regions spanning residues 15–24 and 26–43, followed by a flexible and intrinsically disordered region (**Fig. S1**). Intrinsically disordered proteins as well as numerous miPs interact with macromolecular protein complexes to regulate their function.^29^ To determine whether this was the case for miP-PSTPIP2, a custom antibody was generated and validated^19^ and the endogenous miP was immunoprecipitated from lysates of IL-1β-treated human endothelial cells. Proteins associated with miP-PSTPIP2 were identified by mass-spectrometry-based proteomics, revealing a network of interacting partners with diverse subcellular localizations and functions (**Fig. 1A-B, Dataset S1**). Consistent with its nuclear localization, miP-PSTPIP2 interacted with a large number of nuclear proteins, including members of the BRCA1-A1 and THO complexes, the latter being part of the transcription export machinery. The second largest group of miP-interactors were cytoskeleton proteins, including DAAM1, ARAP3, KIF20A, RHOB and Actin, all of which are involved in actin cytoskeleton remodeling, microtubule-based transport and Rho GTPase signaling. The interaction between miP-PSTPIP2 and actin as well as DAAM1 was validated by immunofluorescence and co-immunoprecipitation (**Fig. 1C-D**). In addition, gene ontology term enrichment analysis revealed a highly significant enrichment in proteins related to caveolar function; including caveolin (CAV)1, CAV2, and caveolae-associated protein (CAVIN)-1-3. Proteins involved in clathrin-mediated intracellular transport, such as the adaptor proteins AP2A2 and AP2B1 were also detected by mass spectrometry and the interactions between miP-PSTPIP2 and CAV1, clathrin heavy chain (CHC) or AP2 were validated by immunofluorescence, proximity ligation assays or co-immunoprecipitation (**Fig. 1E-H**).

**Figure 1.**
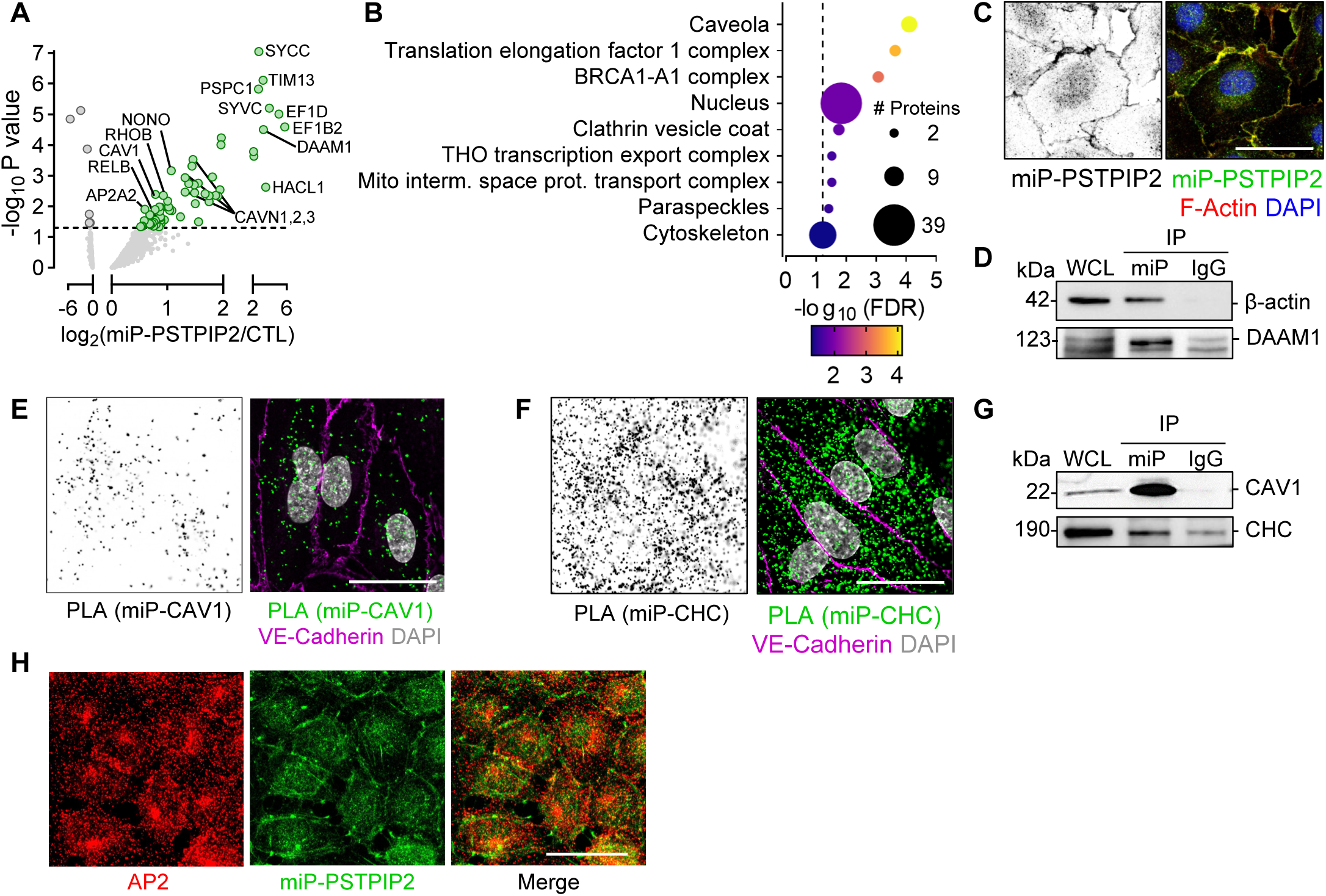
Interactome of miP-PSTPIP2. **A-B,** Volcano plot (A) and GO term enrichment analysis (STRING) (B) of the proteins physically associated with miP-PSTPIP2 (versus rabbit IgGs; CTL) in human endothelial cells treated with IL-1β; n=3 independent cell batches. The dashed lines mark the significance threshold (P value: 0.05). **C,** Representative confocal images showing miP-PSTPIP2 and F-actin in human endothelial cells. Nuclei were stained with DAPI. Scale bar: 10 µm **D,** Co-immunoprecipitation (IP) of miP-PSTPIP2 and β-actin or DAAM1 from human endothelial cell lysates. Similar results were obtained using 2 additional cell batches. **E-F,** Representative confocal images of the interaction (proximity ligation assay: PLA) between miP-PSTPIP2 and caveoilin-1 (CAV1) (E) or Clathrin Heavy Chain (CHC) (F) in human endothelial cells; Scale bar: 10 µm. Similar results were obtained in 3 additional cell batches. **G,** Co-immunoprecipitation (IP) of miP-PSTPIP2 and CAV1 or CHC from human endothelial cell lysates. Similar results were obtained using 3 additional cell batches. **H,** Representative confocal images of AP2A1 and miP-PSTPIP2 in human endothelial cells; Scale bar: 50 µm. Similar results were obtained in 3 additional cell batches.

### miP-PSTPIP2 increases LDL endocytosis and transcytosis

The miP-PSTPIP2 interactome suggested a role for the miP in vesicle trafficking and endocytosis. Therefore, the uptake and intracellular trafficking of transferrin and low-density lipoprotein (LDL) was assessed in human endothelial cells expressing either EGFP (as control) or miP-PSTPIP2. The time-dependent uptake and transcytosis of transferrin were significantly higher in the presence of miP-PSTPIP2 (**Fig. 2A-C**). Similarly, the uptake and transcytosis of LDL, but not oxidized or acetylated LDL, was increased in cells expressing miP-PSTPIP2 (**Fig. 2D-H**). Alterations in endocytic activity are often associated with changes in the abundance of proteins that mediate this process. Therefore, we next examined whether miP-PSTPIP2 affected the expression of several key proteins involved in intracellular trafficking. Overexpression of miP-PSTPIP2 had no effect on the expression of CHC, CAV1, low-density lipoprotein receptor (LDLR), the adaptor protein complex subunits AP2A1 and AP2M1, or the phosphorylation of AP2M1 (**Fig. 3A**). However, consistent with the detected increase in LDL uptake, the membrane expression of LDLR was reduced in the presence of miP-PSTPIP2 (**Fig. 3B**). Next, we examined whether miP-PSTPIP2 localizes with endocytic vesicles during endocytosis by visualizing the localization of the miP and transferrin after incubation with transferrin using total internal reflection fluorescence microscopy. While transferrin-containing endocytic vesicles reached the Golgi apparatus within 15 minutes, miP-PSTPIP2 did not co-localize with transferrin-positive vesicles. In fact, during this process the miP maintained its membrane localization, suggesting that although miP-PSTPIP2 regulates endocytosis it does not do so by does directly associating with the endocytic cargo (**Fig. 3C**).

**Figure 2.**
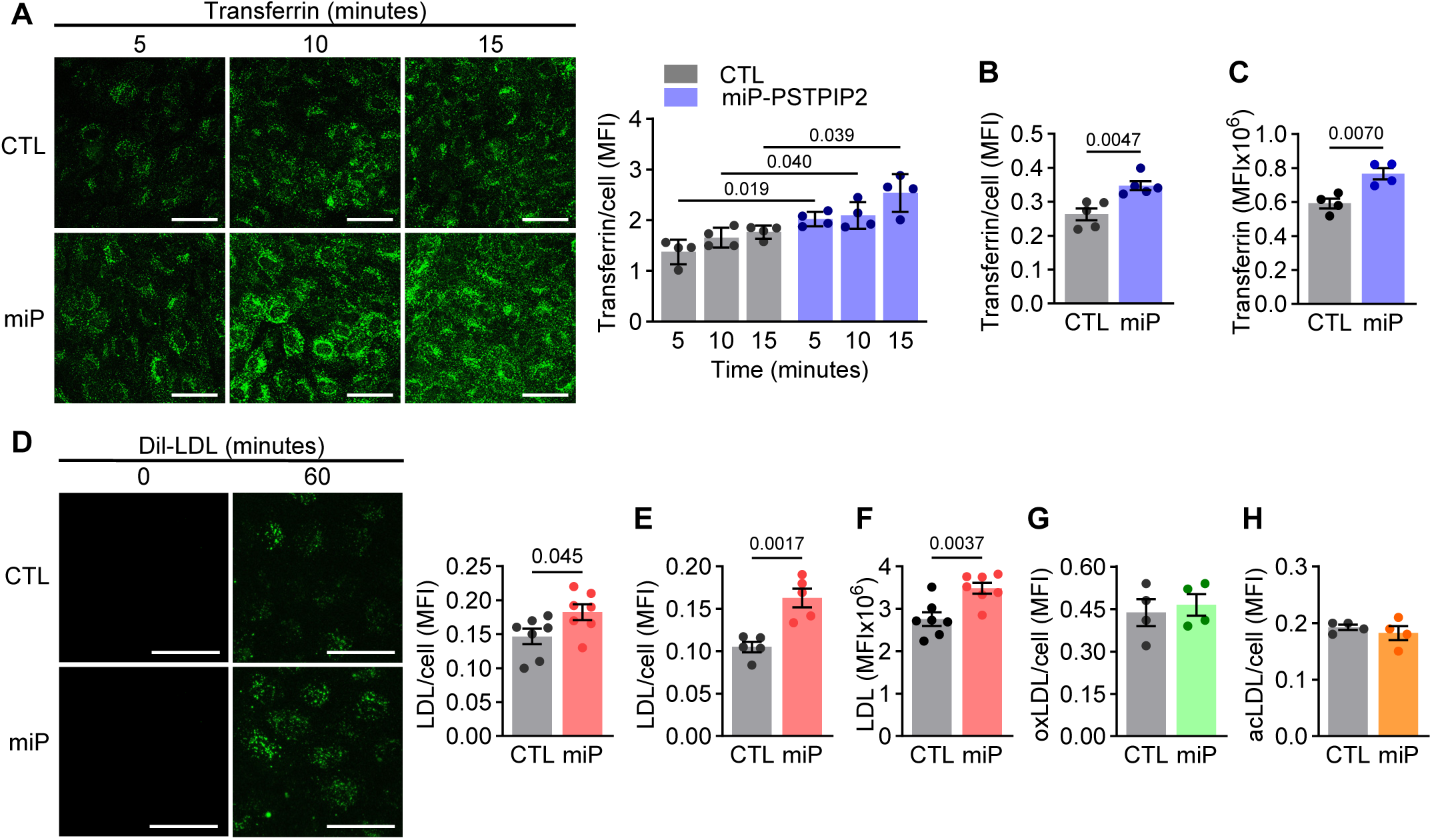
miP-PSTPIP2 modulates endocytosis and transcytosis. **A-B,** Representative confocal images and quantification of transferrin uptake by human endothelial cells (A) or porcine aortic endothelial cells (B) overexpressing miP-PSTPIP2 (miP) or EGFP as control (CTL) after incubation with transferrin for the indicated time; n=4-5 independent cell batches (A: two-way ANOVA and Holm-Šídák’s multiple comparisons test); Scale bar: 50 µm. **C,** Transcytosis of transferrin in human endothelial cells overexpressing miP-PSTPIP2 (miP) or EGFP as control (CTL); n=4 independent cell batches. **D-E,** Representative confocal images and quantification of Dil-LDL uptake by human endothelial cells (D) or porcine aortic endothelial cells (E) overexpressing miP-PSTPIP2 (miP) or EGFP as control (CTL) after incubation with Dil-LDL for 60 minutes; n=5-7 independent cell batches; Scale bar: 50 µm. **F,** Transcytosis of Dil-LDL as in C; n=7 independent cell batches. **G-H,** Uptake (60 minutes) of oxidized LDL (G) or acetylated LDL (H) by human endothelial cells overexpressing miP-PSTPIP2 (miP) or EGFP as control (CTL); n=4 independent cell batches. All statistical analyses were performed using unpaired Student’s t-tests unless otherwise specified.

**Figure 3.**
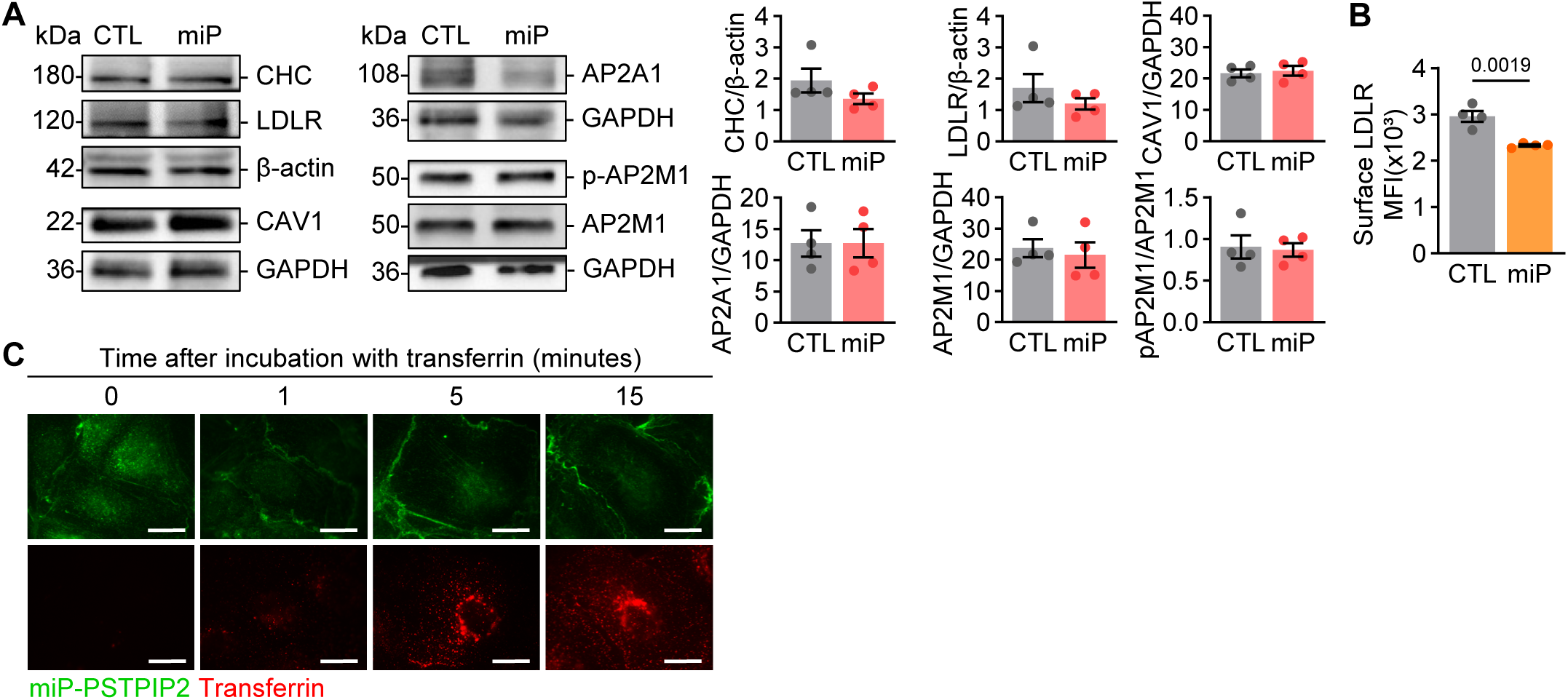
Overexpression of miP-PSTPIP2 has no effect on the expression of proteins involved in endocytosis. **A,** Western blot analysis and quantification of clathrin heavy chain (CHC), low-density lipoprotein receptor (LDLR), Caveolin-1 (CAV1), AP-2 complex subunit alpha-1 (AP2A1), phospho (p-AP2M1) and total AP-2 complex subunit mu (AP2M1) expression in human endothelial cells overexpressing miP-PSTPIP2 (miP) or EGFP as control (CTL). β-Actin or GAPDH were used as loading controls; n=4 independent cell batches. **C,** Flow cytometry analysis of surface LDLR expression in human endothelial cells overexpressing miP-PSTPIP2 (miP) or EGFP (CTL). Data are shown as mean fluorescence intensity (MFI); n=4 independent cell batches. **D,** Representative Total Internal Reflection Fluorescence (TIRF) microscopy images of miP-PSTPIP2 and transferrin in human endothelial cells starved of serum overnight and subsequently incubated with AF555–transferrin for 1, 5 or 15 minutes. n=3 independent experiments; scale bar: 20 µm. All statistical analyses were performed using unpaired Student’s t-tests.

### Effect of miP-PSTPIP2 on cytoskeleton dynamics

Next, we investigated the impact of modulating miP-PSTPIP2 expression on actin cytoskeleton remodeling because the miP associated with actin as well as regulators of cytoskeleton dynamics. To examine the spatial localization of miP-PSTPIP2 in relation to dynamic actin structures, we studied cells treated with cytochalasin D to disrupt the actin cytoskeleton and followed its recovery after washout. MiP-PSTPIP2 was enriched at regions of active actin bundling and at the leading edge of filopodia (**Fig. 4A**), implicating the miP in actin remodeling processes. Consistent with this, endothelial cells expressing miP-PSTPIP2 exhibited accelerated cytoskeleton reorganization and faster restoration of cellular morphology after cytochalasin D treatment compared with control cells (**Fig. 4B**). As the Arp2/3 complex is a key nucleator of branched actin filament networks and plays a key role in lamellipodia formation, vesicle trafficking, and endocytosis,^30,31^ we assessed a possible interaction between miP-PSTPIP2 and this actin remodeling complex. It was possible to demonstrate a physical interaction between miP-PSTPIP2 and Arp3 by proximity ligation assay (**Fig. 4C**). As expected for a mechanism linked to protein interaction, the overexpression of miP-PSTPIP2 did not alter Arp2, Arp3, VASP, RhoA, or cortactin expression (**Fig. 4D**). To test whether these effects were instead mediated through modulation of Arp2/3 complex activity, endothelial cells were treated with CK-666, a selective Arp2/3 inhibitor that blocks nucleation of actin branches. Strikingly, the miP-PSTPIP2–dependent increase in LDL uptake was completely abolished by CK-666 (**Fig. 4E**), demonstrating that miP-PSTPIP2 enhances endocytosis via Arp2/3-driven actin remodeling.

**Figure 4.**
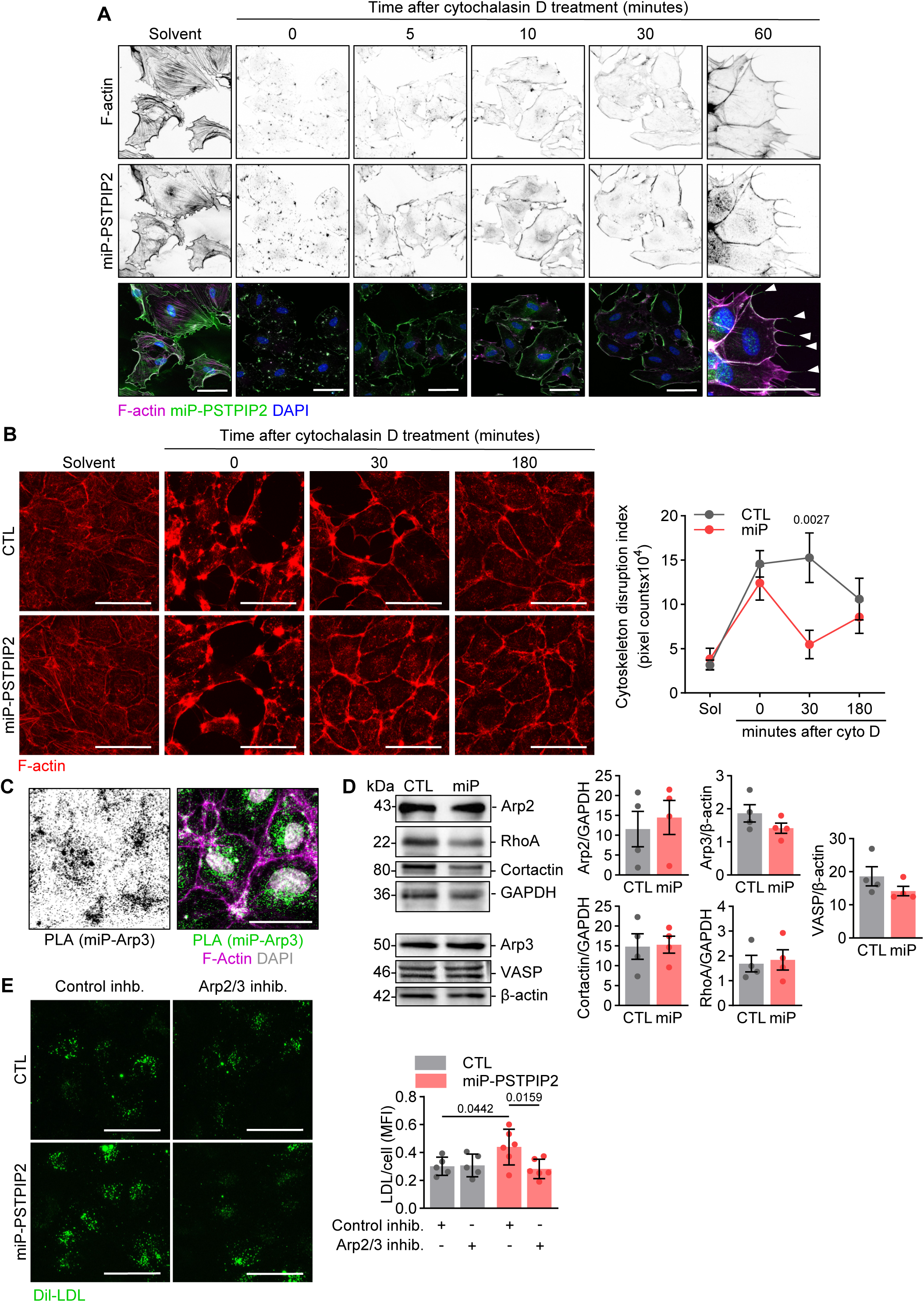
Effect of miP-PSTPIP2 on cytoskeleton dynamics. **A,** Representative confocal images showing miP-PSTPIP2 and F-actin in human endothelial cells under basal conditions (solvent) and 0, 5, 10, 30 or 60 minutes after cytochalasin D treatment (0.5 µmol/L). Similar results were obtained in 2 additional cell batches. Scale bar: 50 µm**. B,** Representative confocal images and quantification of F-actin in human endothelial cells overexpressing miP-PSTPIP2 (miP) or EGFP (CTL) treated with solvent or cytochalasin D (0.5 µmol/L, 20 minutes) for 0, 30 and 180 minutes. The quantification shows the extent of endothelial monolayer disruption measured as counts of low intensity pixels in F-actin immunofluorescence images. n=5 independent cell batches (two-way ANOVA and Holm-Šídák’s multiple comparisons test). Scale bar: 50 µm. **C,** Representative confocal images of the interaction (proximity ligation assay: PLA) between miP-PSTPIP2 and Arp3. Scale bar: 10 µm. Similar results were obtained in 2 additional cell batches. **D,** Western blot analysis and quantification of Actin-related protein 2 (Arp2), Actin-related protein 3 (Arp3), Vasodilator-stimulated phosphoprotein (VASP), WASP-family verprolin homologous protein 1 (WAVE1), Ras homolog family member A (RhoA) and Cortactin expression in human endothelial cells overexpressing miP-PSTPIP2 (miP) or EGFP as control (CTL). β-Actin or GAPDH were used as loading controls; n=4 independent cell batches (unpaired Student’s t-test). **E,** Representative confocal images and quantification of mean fluorescence intensity (MFI) per cell of DiI-LDL uptake in human endothelial cells overexpressing miP-PSTPIP2 (miP) or EGFP (CTL), treated with either control inhibitor CK689 or Arp2/3 inhibitor CK666. n=4 independent cell batches (two-way ANOVA and Holm-Šídák’s multiple comparisons test). Scale bar: 50 µm.

### miP-PSTPIP2 increases vascular permeability and decreases endothelial cell migration

To determine whether miP-PSTPIP2 was implicated in other cellular processes reliant on dynamic actin remodeling, we assessed its impact on endothelial cell migration and barrier function. Endothelial cells that expressed miP-PSTPIP2 demonstrated impaired collective migration in scratch wound assays as well as diminished chemotactic migration towards fetal bovine serum (**Fig. 5A-B**). Consistent with its upregulation by inflammation,^19^ endothelial cell barrier function was impaired by miP-PSTPIP2 as demonstrated by enhanced permeability to 40 kDa dextran (**Fig. 5C**). Similarly, the monocyte chemoattractant protein-1-induced migration of monocytes through endothelial cells was significantly enhanced upon miP-PSTPIP2 expression (**Fig. 5D**).

**Figure 5.**
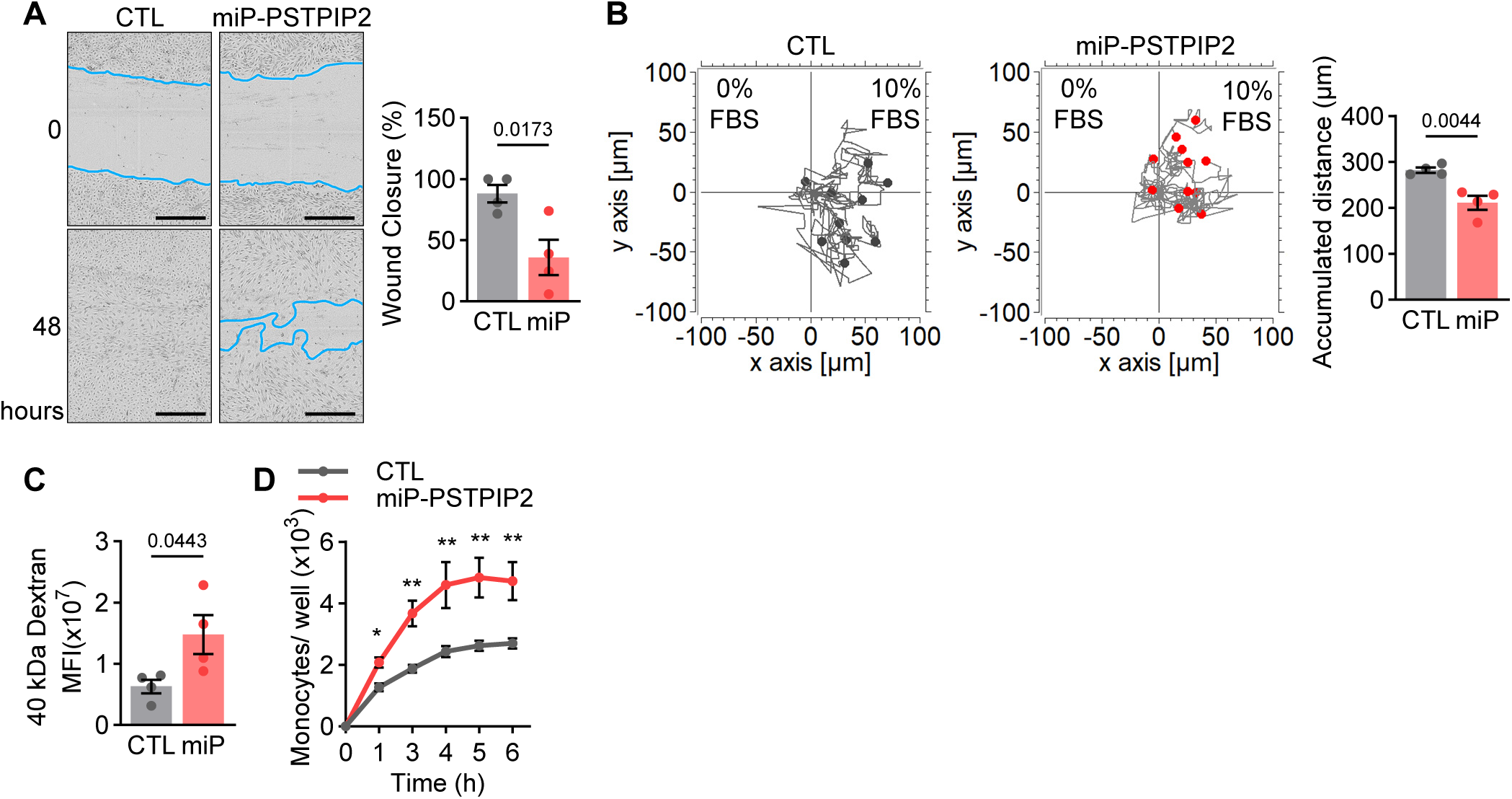
miP-PSTPIP2 decreases cell migration and increases vascular permeability. **A,** Representative images and quantification of human endothelial cells overexpressing miP-PSTPIP2 (miP) or EGFP (CTL) at 0 h and 48 h after scratching; wound edges are highlighted in blue. Scale bar: 500 µm; n=4 independent cell batches. **B,** Migration of human endothelial cells overexpressing miP-PSTPIP2 (miP) or EGFP (CTL) toward 10% FBS. Cell trajectories were recorded over 20 hours. Representative migration tracks from one experimental batch are shown; n=4 independent cell batches. **C,** Permeability of monolayers of human endothelial cell overexpressing miP-PSTPIP2 (miP) or EGFP (CTL) to fluorescently-labelled 40 kDa dextran. Mean fluorescence intensity (MFI) in the lower chamber of the transwell was measured after incubation for 15 minutes; n=4 independent cell batches. **D,** Monocyte chemotactic protein-1-induced migration of monocytes through human endothelial cell monolayers overexpressing miP-PSTPIP2 (miP) or EGFP (CTL); n=4 independent cell batches (two-way ANOVA and Holm-Šídák’s multiple comparisons test). All other statistical analyses were performed using unpaired Student’s t-tests.

## DISCUSSION

This study identifies miP-PSTPIP2 as a novel regulator of endothelial cytoskeleton dynamics and vesicular trafficking. miP-PSTPIP2 interacts with actin-associated and endocytic proteins, enhances LDL and transferrin uptake through Arp2/3-dependent actin remodeling and increases endothelial permeability while impairing directed migration. These findings suggest that miP-PSTPIP2 links inflammatory signaling to cytoskeleton reorganization and vascular dysfunction.

MiPs often exert regulatory effects through binding to larger protein complexes or membranes.^1,32^ We found that miP-PSTPIP2 associates with the cytoskeleton and interacts with proteins involved in endocytic machinery and actin remodeling, suggesting a role in vesicular trafficking. Caveolae and clathrin-coated pits are the primary endocytic platforms in endothelial cells and control the internalization of extracellular molecules and receptors such as transferrin and LDL.^33,34^ Functionally, overexpression of miP-PSTPIP2 selectively enhanced receptor-mediated endocytosis and transcytosis of LDL and transferrin without affecting scavenger-mediated uptake, suggesting that its role is specific to physiological endocytic pathways rather than generalized membrane internalization. This selective effect is particularly relevant for atherogenesis, as native LDL uptake contributes to physiological lipid handling, whereas modified LDL species such as oxLDL promote foam cell formation and vascular inflammation.

Despite its clear effect on endocytosis and transferrin uptake, miP-PSTPIP2 did not associate with the endocytic machinery or intracellular vesicles during the endocytic process, nor did it alter AP2 subunit expression or AP2M1 phosphorylation, which are critical for clathrin-coated vesicle formation.^35^ These findings suggest that miP-PSTPIP2 likely acts at the level of the plasma membrane or during vesicle formation. We hypothesized that the effects of miP-PSTPIP2 may be mediated through actin cytoskeleton remodeling, which is known to facilitate endocytosis by generating cortical tension, driving membrane invagination, and supporting vesicle scission.^36,37,29,30^ Consistent with this model, miP-PSTPIP2 localized to regions of active actin remodeling, including the tips of filopodia, cell-cell junctions, and actin bundles. Upon actin disruption and reorganization, cells overexpressing miP-PSTPIP2 regained their morphology faster and nucleated actin filaments more efficiently. These findings suggest that miP-PSTPIP2 enhances endocytosis by modulating cortical actin dynamics, which are critical for membrane invagination, thereby enhancing the formation of endocytic vesicles.^38,39,37^ The Arp2/3 complex, along with nucleation-promoting factors such as VASP, RhoA, and cortactin, is a key regulator of the generation of branched actin networks necessary for both lamellipodia formation and endocytic vesicle budding.^30,40^ Notably, we found that pharmacological inhibition of the Arp2/3 complex abolished the miP-dependent increase in LDL uptake, confirming that miP-PSTPIP2 enhances endocytosis in an Arp2/3-dependent manner.

The functional consequences of miP-PSTPIP2 on cytoskeleton remodeling extend beyond endocytosis. Actin dynamics are critical for endothelial migration and barrier function, particularly under conditions of vascular stress or injury. Consistent with this, miP-PSTPIP2 overexpression impaired wound closure and directed migration while increasing vascular permeability and monocyte transmigration, processes highly relevant to atherogenesis.^41,42^ Emerging evidence indicates that excessive or misdirected Arp2/3-activity can compromise effective migration, as over-branched cytoskeleton networks or misallocated actin polymerization hinder lamellipodia extension and directional movement.^43,31^ These observations suggest that miP-PSTPIP2 may redirect Arp2/3 activity preferentially toward endocytic and nucleation events, enhancing vesicle internalization and cortical remodeling at the expense of coordinated migration. Beyond its role in endocytic trafficking and migration, miP-PSTPIP2 modulated endothelial barrier function. Overexpression of the miP increased dextran permeability and monocyte transmigration, consistent with compromised cell-cell junctions. Given its localization to regions of active actin remodeling and its enhancement of Arp2/3 activity, these effects likely arise from altered actin dynamics at junctional sites. By shifting the balance to cortical actin polymerization, miP-PSTPIP2 may destabilize adherens and tight junctions, thereby increasing paracellular permeability.

In summary, miP-PSTPIP2 modulates actin cytoskeleton remodeling, selective endocytosis, and endothelial barrier function. Prolonged or excessive miP-PSTPIP2 expression may contribute to dysregulated lipid handling, pathological permeability and impaired endothelial cell-mediated regeneration, which are hallmarks of chronic inflammation leading to vascular disease. These findings highlight the broader functional significance of endothelial cell miPs and underscore the potential for their targeting in the treatment of cardiovascular disease.

## ACKNOWLEDGEMENTS

The authors thank Sandrine Ngaha, Isabel Winter and Mechtild Piepenbrock-Gyamfi for expert technical assistance.

---

**Supplementary Figure 1.**
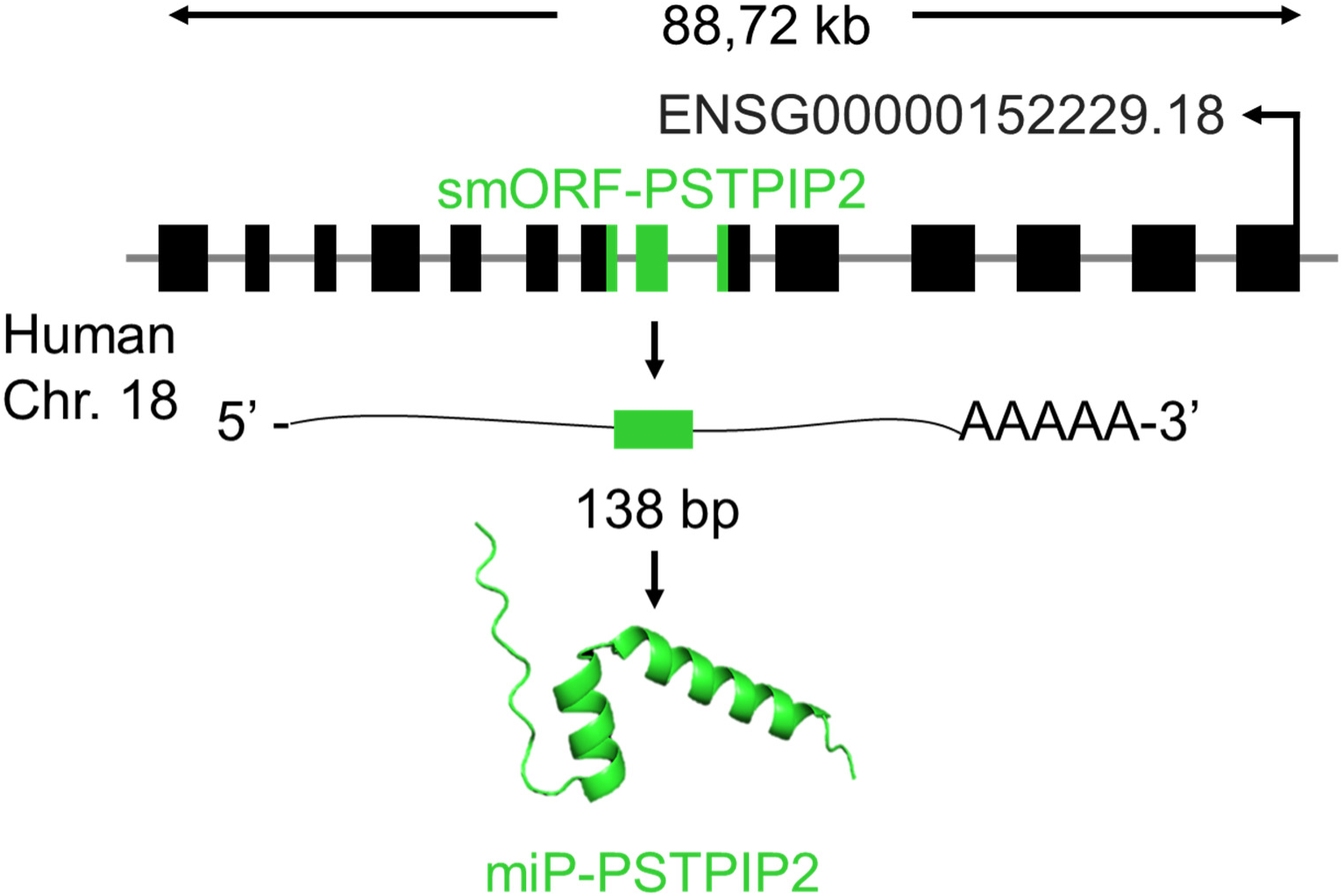
Genomic location of smORF-PSTPIP2 and structural prediction of miP-PSTPIP2. Location of smORF-PSTPIP2 (green) in the *PSTPIP2* gene and transcript and secondary structure prediction of miP-PSTPIP2 generated with AlphaFold3.

